# Mechanism and role of astrogliosis in the pathogenesis of HIV-associated pain One Sentence Summary: Neuron-to-astrocyte Wnt5a signaling-activated astrogliosis is essential for the development of pathological pain induced by HIV-1 gp120

**DOI:** 10.1101/2021.04.28.441838

**Authors:** Xin Liu, Chilman Bae, Benjamin Gelman, Jin Mo Chung, Shao-Jun Tang

## Abstract

Pathological pain is the most common neurological disorder in people living with HIV-1/AIDS (PLWHA), and rationale-based effective treatment is not available. Multiple neuropathologies develop in the pain transmission pathways in of HIV patients, consistent with their nociceptive dysfunction^1,2^. One of the prominent neuropathologies associating with the manifestation of pain in HIV patients is astrogliosis (a.k.a. reactive astrocytes) in the spinal dorsal horn (SDH)^1^, the spinal center for the transmission of pain signals from peripheral organs to the brain. However, the pathogenic role and the activation mechanism of astrogliosis are unclear. Here, we show that the astrogliosis is crucial for the pain pathogenesis induced by HIV-1 gp120, a key etiologically relevant protein^2^, and that a neuron-to-astrocyte Wnt5a signal controls the astrogliosis. We found that ablation of astrogliosis blocked the development of gp120-induced mechanical hyperalgesia, and concomitantly the expression of neural circuit polarization (NCP) in the SDH. In addition, we demonstrated that conditional knockout (CKO) of either Wnt5a in neurons or its receptor ROR2 in astrocytes abolished not only gp120-induced astrogliosis but also the hyperalgesia and the NCP. Furthermore, we found that the astrogliosis promoted expression of the NCP and the hyperalgesia via IL-1β regulated by a Wnt5a-ROR2-MMP2 axis. Our results elucidate an important role and a novel mechanism of astrogliosis in the pathogenesis of HIV-associated pain. Targeting reactive astrocytes by manipulating the mechanistic processes identified here may lead to the development of effective therapy to treat the pain syndrome in HIV patients.

## INTRODUCTION

Among over 38 million people living with HIV-1/AIDS (PLWHA) worldwide, more than 50% develop chronic pain^3,4^. Even in post-cART (combinatorial anti-retroviral therapy) era, HIV-associated pain remains highly prevalent^3,5^, continues to have a major impact on the quality of life of the patients^6^, and is still a central clinical and public health concern^4,7^. The HIV-associated pain syndrome is manifested in various forms, including peripheral neuropathy, headache and visceral pain, and is often undertreated. The current treatments by opioid or non-opioid analgesics are symptomatic management adapted from pain control used in other diseases, and have limited effectiveness^8,9^. Development of rational-based effective pain therapy for HIV patients is an urgent unmet need. Towards this goal, it is essential to elucidate the underlying pathogenic mechanisms, which is still poorly understood.

Several etiological factors have been considered in pain pathogenesis in PLWHA, including HIV-1 virus and proteins, opportunistic infection and antiretroviral therapy. Early studies indicate a correlation between HIV-1 viral load and pain development. However, pain remains highly prevalent in the post cART era^3,5^, indicating viral replication and infection suppressed by antiretroviral drugs are not the major cause responsible for the pain pathogenesis^2^. Both HIV-1 neurotoxic proteins and cART are thought to play crucial roles in the pain pathogenesis. The expression of individual HIV proteins from defective proviruses is observed even under the cART treatment^2,10,11^. A critical pathogenic contribution of HIV-1 gp120 protein is suggested by its high level in the postmortem SDH from HIV patients who developed pain but not from the patients who did not manifest pain^2^, and its activity in inducing HIV pain-related pathologies in animal models^2,12–15^. Tat and vpr proteins also induce pain in animals^16,17^. However, unlike gp120, their expression levels in the SDH do not differentiate pain-positive from pain-negative HIV patients^2^. Anti-retroviral drugs, especially nucleoside reverse transcriptase inhibitors (NRTIs), may also contribute to the pain pathogenesis of HIV patients on cART^18,19^. Both HIV-1 protein and NRTIs may elicit neuroinflammation that cause neuronal damages in the pain circuits^2,12,15,18,20^.

Neuronal pathological changes develop in both the peripheral and the central pain processing pathways of HIV patients who developed chronic pain. In the peripheral nervous system (PNS), painful sensory neuropathy, manifested as dying-back degeneration of the peripheral endings of the sensory fibers, is a common neuropathology, especially in the limbs^21^. Degenerating pain sensory neurons may become spontaneously activated, or sensitized to normally non-noxious stimuli. In the central nervous system (CNS), there is clear synapse degeneration specifically in the SDH of pain-positive HIV patients^2^. Loss of synapses, especially inhibitory synapses, may lead to central sensitization by causing excitatory/inhibitory imbalance of neural circuits. Both peripheral neuropathy and synapse degeneration are simulated in the mouse pain models induced by gp120^2,22^.

Neuroinflammation is another hallmark in the SDH of pain-positive HIV patients, as manifested by the up-regulation of proinflammatory mediators and the activation of astroctyes^1,2^. Astrocytes are the most abundant glia, and play essential roles in regulating CNS homeostasis^23^ and plasticity of neural circuits^24^, including synaptogenesis, synapse pruning, synaptic transmission and plasticity^25–27^. Under neurological disease conditions such as neurodegeneration and pathological pain^28–30^, astroglia often become activated and secret inflammatory and other factors that may modulate pathogenesis. The possible involvement of astrogliosis in pain pathogenesis was suggested^29,31,32^. The prominent astroglial activation in the SDH specifically in PLWHA who develop pain suggests reactive astrocytes play a critical role in the pathogenesis of HIV-associated pain. However, the mechanism that triggers astrogliosis and the specific pathogenic role that astrogliosis plays remain unclear.

In this work, we use the mouse models to determine the role and activation mechanisms of astrogliosis during the pain pathogenesis induced by gp120. The results show that astrogliosis is essential for gp120 to induce mechanical allodynia and polarization of pain neural circuits in the SDH, and that Wnt5a-ROR2 is a neuron-to-astrocyte signaling pathway that controls the gp120-induced astrogliosis. These findings elucidate key molecular and cellular mechanisms of HIV-associated pain pathogenesis, and may help design effective pain therapy for HIV patients.

## RESULTS

### Astrogliosis is crucial for HIV-1 gp120 to induce pathological pain

We used an established mouse model of HIV-associated pain, generated by intrathecal (i.t.) injection of gp120 (4μg/kg, i.t., daily at the first 2 days and then every other day, 4 times), which extensively simulates observed pain-related pathologies in pain-positive HIV patients, including neuropathy, synapse degeneration, and astrogliosis^2^. To directly test the role of astrogliosis in the development of gp120-induced pain, we employed a genetic approach to selectively ablate proliferating reactive astrocytes, using the GFAP-thymidine kinase (TK) transgenic mice^33^, which can express gp120-induced pain similarly as WT mice (Fig.1B). We intrathecally (i.t.) injected ganciclovir (GCV; 5mg/kg) at the first two days after gp120 administration to ablate astrogliosis. The GCV treatment blocked the gp120-induced increase of GFAP in the transgenic but not in WT mice (Fig. 1A), indicating specific ablation of reactive astrocytes. Importantly, von Frey tests showed that GCV blocked the expression of gp120-induced mechanical hyperalgesia in GFAP-TK but not in WT mice (Fig. 1B). These results suggest that astrogliosis plays a critical role in the pathogenesis of gp120-induced pain.

**Figure 1.**
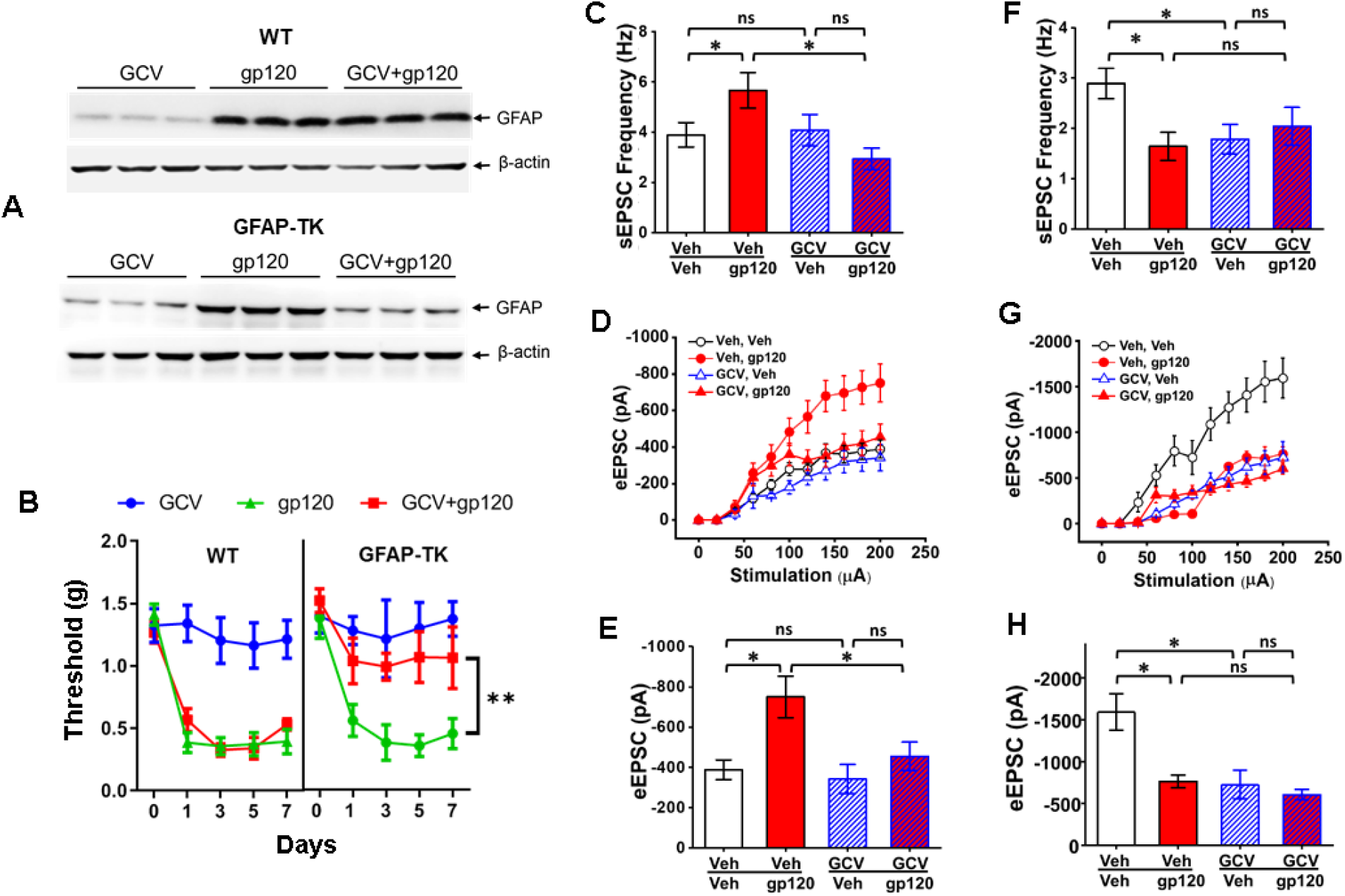
Ablation of astrogliosis blocks gp120-induced hyperalgesia and neural circuit polarization (NCP) in the spinal dorsal horn (SDH). **A.** Immunoblotting analysis of spinal GFAP proteins. Ganciclovir (GCV) administration (5mg/kg, daily, first 2 days, i.t.) ablated astrogliosis induced by gp120 (4μg/kg, daily at the first 2 days and then every other day, four times, i.t.) in GFAP-TK transgenic but not in wild-type (WT) mice. Tissues were collected at Day 7 and GFAP protein levels were determined by immunoblotting analysis. **B.** Von Frey tests. GCV administration blocked the expression of gp120-induced mechanical hyperalgesia in the GFAP-TK transgenic but not WT mice (5 mice/group; **, p<0.01). **C.** Effect of ablation of astrogliosis on spontaneous EPSCs (sEPSCs) of non-tonic firing neurons in the SDH of GFAP-TK transgenic mice. Whole cell patch recording was performed on spinal slices prepared from GFAP-TK transgenic mice treated with gp120 and/or GCV (at Day 7 in B). gp120 significantly increased sEPSC frequency, and GCV abolished the increase. GCV did not affect the basal level of sEPSC frequency. Veh/Veh: 21/3 (cells/animals); veh/gp120: 20/3; GCV/Veh: 29/4; GCV/gp120: 47/4. **D**. Effect of astrogliosis ablation on evoked EPSCs (eEPSCs) of non-tonic firing neurons in the SDH of GFAP-TK transgenic mice. eEPSC amplitudes of patched SDH neurons were recorded on spinal slices prepared as in C. gp120 increased eEPSC amplitudes, and GCV abolished the increase. GCV did not affect the basal level of eEPSC amplitudes. Veh/Veh: 24/3 (cells/animals); veh/gp120: 24/4; GCV/Veh: 35/3; GCV/gp120: 24/4. **E**. Statistical analysis of eEPSC amplitudes evoked by 200μA stimulation shown in D. **F**. Effect of astrogliosis ablation on sEPSCs of tonic firing neurons in the SDH of GFAP-TK transgenic mice. gp120 significantly decreased sEPSC frequency. Although GCV treatment by itself also decreased sEPSC frequency, gp120 did not cause further decrease after GCV treatment. Veh/Veh: 13/3 (cells/animals); veh/gp120: 10/3; GCV/Veh: 20/3; GCV/gp120: 26/4. **G**. Effect of astrogliosis ablation on eEPSCs of tonic firing neurons in the SDH of GFAP-TK transgenic mice. gp120 decreased eEPSC amplitudes. GCV also decreased eEPSC amplitudes by itself, but gp120 did not further decrease the amplitudes after GCV administration. Veh/Veh: 14/3 (cells/animals); veh/gp120: 12/4; GCV/Veh: 20/3; GCV/gp120: 16/4. **H**. Statistical analysis of eEPSC amplitudes evoked by 200μA stimulation shown in G. *, p<0.05; ns, p>0.05.

Gp120 also caused microglial activation (Sup. Fig. 1). To test the potential role of microglia in gp120-induced pain, we ablated microglia with colony-stimulating factor 1 receptor (CSF1R) inhibitor PLX5622 as reported^34^ (Sup. Fig. 1A). We observed that microglial ablation only partially inhibited the early expression of the gp120-induced mechanical hyperalgesia, without affecting the late stage of pain development and maintenance (Sup. Fig. 1B). These results indicate that microglial activation only modulates the development of gp120-induced pain at its early stage. This interpretation is consistent with the prior observation that microglial activation is not evident in the postmortem spinal tissues from pain-positive HIV patients who may have established pain states long before they died^1^.

### Gp120 disturbs homeostasis of pain neural circuitry in the SDH via astrogliosis

Next, we sought to understand the neural circuit pathogenesis underlying gp120-induced pain and the contribution of the astrogliosis to the circuit pathogenesis. Whole-cell recording of SDH neurons in GFAP-TK mice showed that gp120 increased both the frequency of spontaneous excitatory postsynaptic currents (sEPSCs) and the amplitude of evoked EPSCs (eEPSCs) of excitatory neurons (Fig. 1C-E), identified by their characteristic non-tonic firing patterns^35,36^. By contrast, gp120 decreased both the sEPSC frequency and eEPSC amplitude of inhibitory neurons (Fig. 1F-H), identified by their characteristic tonic firing pattern^35,36^. These results suggest that gp120 causes increase of excitatory inputs to excitatory neurons and decrease of excitatory inputs to inhibitory neurons. This gp120-induced neural circuit polarization (NCP) may facilitate activation of the SDH pain pathway and consequent hyperalgesia.

How might astrogliosis contribute to gp120-induced pathogenesis in the pain neural circuit? To answer this question, we tested the effect of astrogliosis ablation on gp120-induced NCP in the SDH of GFAP-TK mice. We observed that astrogliosis ablation by GCV abolished the gp120-induced increase in sEPSC frequency and eEPSC amplitude of excitatory neurons (Fig. 1C-E). We found that GCV by itself caused decrease of sEPSC frequency and eEPSC amplitude of inhibitory neurons; nonetheless, it prevented further gp120-induced decreases in sEPSC frequency and eEPSC amplitude (Fig. 1F-H). Together, these results indicate that astrogliosis is critical for NCP expression induced by gp120.

### Neuronal Wnt5a is critical for gp120-induced astrogliosis, pain and NCP

Wnt5a is a secreted signaling protein mainly expressed by neurons^37,38^. Its secretion is regulated by synaptic activity^39,40^. Wnt5a is upregulated in the SDH of pain-positive HIV patients^38^ and the gp120 pain model^37,41^, and is implicated in pain pathogenesis^42–44^. Wnt5a antagonist impairs astrogliosis in pain models^18,45^. We hypothesized neuronal Wnt5a played a critical role in gp120-induced astrogliosis. To test this hypothesis, we generated neuronal Wnt5a conditional knock-out (CKO) mice by crossing floxed Wnt5a mice^46^ with synapsin 1-Cre mice^47^. We found that the Wnt5a CKO blocked gp120-induced Wnt5a up-regulation (Fig. 2A), indicating that neurons were the source of Wnt5a induced by gp120. Importantly, the Wnt5a CKO abolished gp120-induced astrogliosis (Fig. 2B), but not microglial activation (Sup. Fig. 2). These data show that neuronal Wnt5a is crucial for gp120-induced astrogliosis.

**Figure 2.**
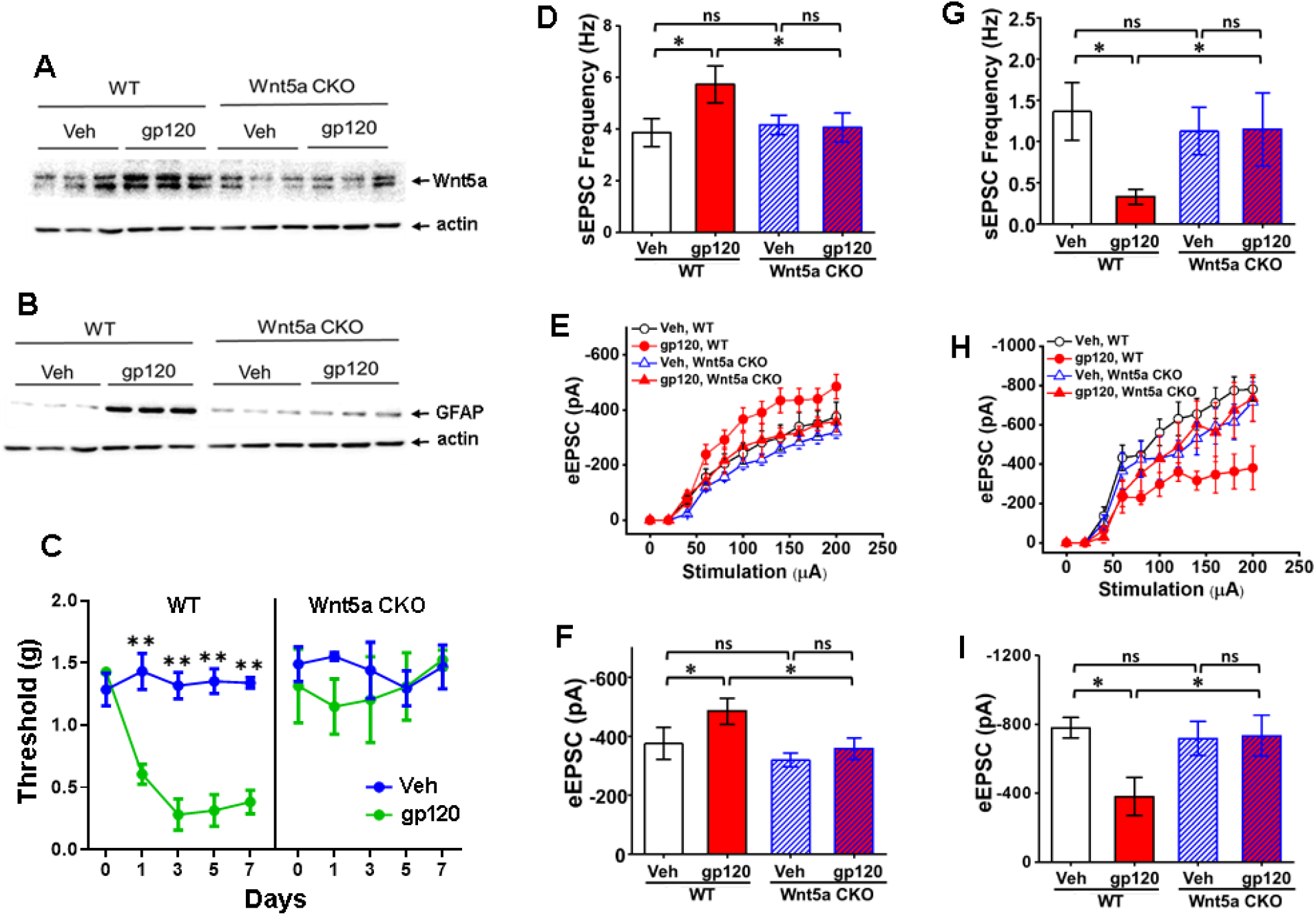
Wnt5a CKO in neurons abolishes gp120-induced astrogliosis, hyperalgesia and NCP. **A**. Immunoblotting analysis showed that neuronal Wnt5a CKO abolished Wnt5a up-regulation induced by gp120. Tissues were collected at Day 7. **B**. Immunoblotting analysis of spinal GFAP proteins indicated that neuronal Wnt5a CKO blocked astrogliosis induced by gp120 (described in Fig 1A legend) Tissues were collected at Day 7. **C.** Von Frey tests showed that neuronal Wnt5a CKO blocked the expression of gp120-induced mechanical hyperalgesia (5 mice/group; **, p<0.01). **D.** Neuronal Wnt5a CKO blocked gp120-induced increase of frequency of spontaneous EPSCs (sEPSCs) of non-tonic firing neurons in the SDH. Whole cell patch recording was performed on spinal slices as in Fig. 1. gp120 significantly increased sEPSC frequency in WT but not in the Wnt5a CKO mutant mice. Veh/WT: 19/3 (cells/animals); WT/gp120: 21/4; Wnt5a CKO/Veh: 57/5; Wnt5a CKO/gp120: 40/4. **E**. Effect of neuronal Wnt5a CKO on evoked EPSCs (eEPSCs) of non-tonic firing neurons in the SDH. eEPSC amplitudes were recorded on spinal slices prepared as in D. gp120 increased eEPSC amplitudes in WT but not in the Wnt5a CKO mice. Veh/WT: 24/3 (cells/animals); WT/gp120: 24/4; Wnt5a CKO/Veh: 55/5; Wnt5a CKO/gp120: 53/4. **F**. Statistical analysis of eEPSC amplitudes evoked by 200μA stimulation shown in E. **G**. Effect of the neuronal Wnt5a CKO on sEPSCs of tonic firing neurons in the SDH. gp120 significantly decreased sEPSC frequency in WT but not in the Wnt5a CKO mutant mice. Veh/WT: 12/3 (cells/animals); WT/gp120: 15/4; Wnt5a CKO/Veh: 28/5; Wnt5a CKO/gp120: 9/4. **H**. Effect of neuronal Wnt5a CKO on eEPSCs of tonic-firing neurons in the SDH. gp120 decreased eEPSC amplitudes in the WT but not in the Wnt5a CKO mutant mice. Veh/WT: 8/3 (cells/animals); WT/gp120: 10/3; Wnt5a CKO/Veh: 24/3; Wnt5a CKO/gp120: 10/3. **I**. Statistical analysis of eEPSC amplitudes evoked by 200μA stimulation shown in H. *, p<0.05; ns, p>0.05.

In addition to blockade of the astrogliosis, the neuronal Wnt5a CKO also abolished the expression of gp120-induced mechanical hyperalgesia (Fig. 2C). Furthermore, the Wnt5a CKO blocked the gp120-induced increase in sEPSC frequency and eEPSC amplitude of excitatory neurons (Fig. 2D-2F) and the decrease in sEPSC frequency and eEPSC amplitude of inhibitory neurons (Fig. 2G-2I). These findings indicate that neuronal Wnt5a mediates the expression of gp120-induced mechanical hyperalgesia and NCP. Because astrogliosis is essential for the gp120-induced hyperalgesia and NCP (Fig. 1), the inhibitory effects of the Wnt5a CKO on gp120-induced astrogliosis, hyperalgesia and NCP (Fig. 2) suggest that Wnt5a regulates gp120-induced hyperalgesia and NCP by activating astrogliosis.

### Gp120-induced astrogliosis, pain and NCP depend on astrocytic ROR2 receptor

What receptor on astrocytes would mediate Wnt5a’s activity in stimulating astrogliosis? To address this problem, we examined the role of ROR2, a Wnt5a co-receptor that is expressed in astrocytes in the SDH^37^. Astrocytic ROR2 CKO mice were generated by crossing the floxed ROR2^48^ and GFAP-Cre^49^ mouse lines. We found that the ROR2 CKO also blocked gp120-induced astrogliosis (Fig. 3A). The astrogliosis blockade by the ROR2 CKO was incomplete but significant (Fig. 3A). By contrast, the blockade effect of neuronal Wnt5a CKO on gp120-induced astrogliosis was more complete (Fig. 2). These observations indicate that ROR2 is not the sole pathway through which Wnt5a activates astrocytes.

**Figure 3.**
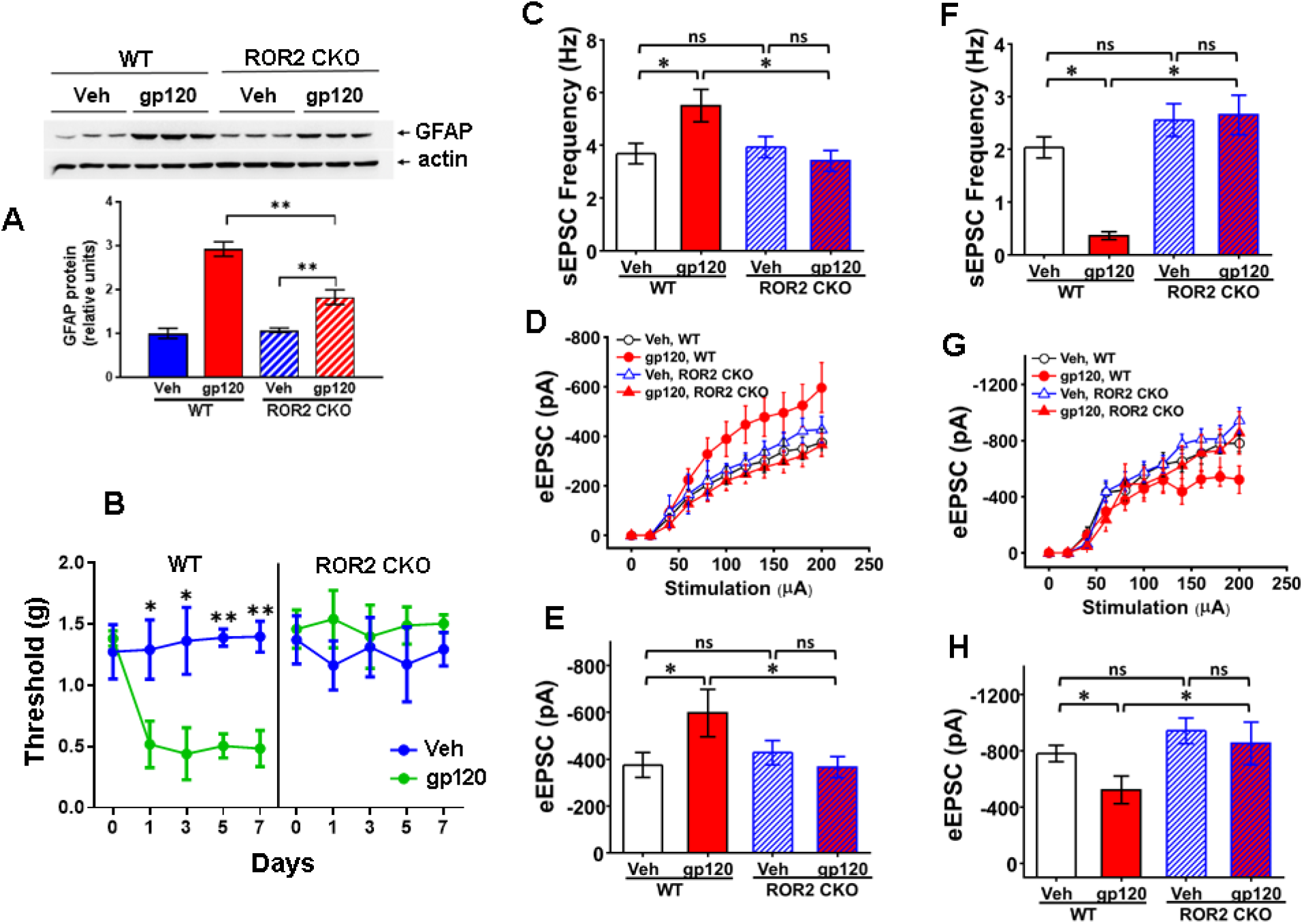
Astrocytic ROR2 CKO in diminishes gp120-induced astrogliosis, hyperalgesia and NCP. **A**. Immunoblotting analysis OF spinal GFAP protein showed that astrocytic ROR2 CKO significantly impaired astrogliosis induced by gp120 (described in Fig 1A legend). Tissues were collected at Day 7 **B.** Von Frey tests showed that astrocytic ROR2 CKO inhibited the expression of gp120-induced mechanical hyperalgesia. 5 mice/group; *, p<0.05; **, p<0.01. **C.** Astrocytic ROR2 CKO inhibited gp120-induced increase of sEPSC frequency of non-tonic firing neurons in the SDH. Whole cell patch recording was performed on spinal slices as in Fig. 1. gp120 significantly increased sEPSC frequency in WT but not in the ROR2 CKO mutant mice. Veh/WT: 32/3 (cells/animals); WT/gp120: 18/5; ROR2 CKO/Veh: 45/3; ROR2 CKO/gp120: 32/4. **D**. Effect of astrocytic ROR2 CKO on eEPSCs of non-tonic firing neurons in the SDH. gp120 increased eEPSC amplitudes in WT but not in the ROR2 CKO mice. Veh/WT: 24/3 (cells/animals); WT/gp120: 36/4; ROR2 CKO/Veh: 34/3; ROR2 CKO/gp120: 38/3. **E**. Statistical analysis of eEPSC amplitudes evoked by 200μA stimulation shown in D. **F**. Effect of the astrocytic ROR2 CKO on sEPSCs of tonic firing neurons in the SDH. gp120 significantly decreased sEPSC frequency in WT but not in the ROR2 CKO mutant mice. Veh/WT: 19/3 (cells/animals); WT/gp120: 24/4; ROR2 CKO/Veh: 22/3; ROR2 CKO/gp120: 31/3. **G**. Effect of astrocytic ROR2 CKO on eEPSCs of tonic-firing neurons in the SDH. gp120 decreased eEPSC amplitudes in the WT but not in the ROR2 CKO mutant mice. Veh/WT: 8/3 (cells/animals); WT/gp120: 11/3; ROR2 CKO/Veh: 16/3; ROR2 CKO/gp120: 18/3. **H**. Statistical analysis of eEPSC amplitudes evoked by 200μA stimulation shown in H. *, p<0.05; ns, p>0.05.

Similar to the effect of the neuronal Wnt5a CKO (Fig. 2), the astrocytic ROR2 CKO also abolished gp120-induced mechanical hyperalgesia (Fig. 3B). Furthermore, the astrocytic ROR2 CKO also blocked gp120-induced increase in sEPSC frequency and eEPSC amplitude of excitatory neurons (Fig. 3C-3E) and the decrease in sEPSC frequency and eEPSC amplitude of inhibitory neurons in the SDH (Fig. 3F-3H). These findings suggest that astrocytic ROR2 simulates the effect of the neuronal Wnt5a CKO in regulating gp120-induced astrogliosis, pain, and NCP.

### Wnt5a/ROR2-regulated astrogliosis promotes gp120-induced pain pathogenesis via IL-1β

The effects of the Wnt5a CKO and the ROR2 CKO on gp120-induced astrogliosis described above suggest a neuron-to-astrocyte Wnt5a-ROR2 signaling pathway that activates astrocytes. How would the Wnt5a signaling-dependent astrogliosis regulate gp120-induced pain pathogenesis? Prior studies indicate that reactive astrocytes might regulate pain pathogenesis by releasing proinflammatory mediators such as IL-1β^29^. We observed IL-1β expression in astrocytes in the SDH of the gp120-treated mice (Fig. 4A). gp120 up-regulates IL-1β via Wnt5a signaling^37^. Using the GFAP-TK mouse, we further showed that the gp120-induced up-regulation of IL-1β was blocked by ablating astrogliosis (Fig. 4B), confirming reactive astrocytes as the source of gp120-induced IL-1β suggested by the immunostaining results (Fig. 4A). Similar to its effect on astrogliosis induced by gp120 (Fig. 2B), the neuronal Wnt5a CKO abolished gp120-induced IL-1β upregulation (Fig. 4C). The astrocytic ROR2 CKO significantly but not completely inhibited the IL-1β upregulation (Fig. 4D), similar to its incomplete inhibition on the astrogliosis (Fig. 3A). These results show that neuron-to-astrocyte Wnt5a signaling controls gp120-induced IL-1β upregulation, at least partially via ROR2-regulated reactive astrocytes.

**Figure 4.**
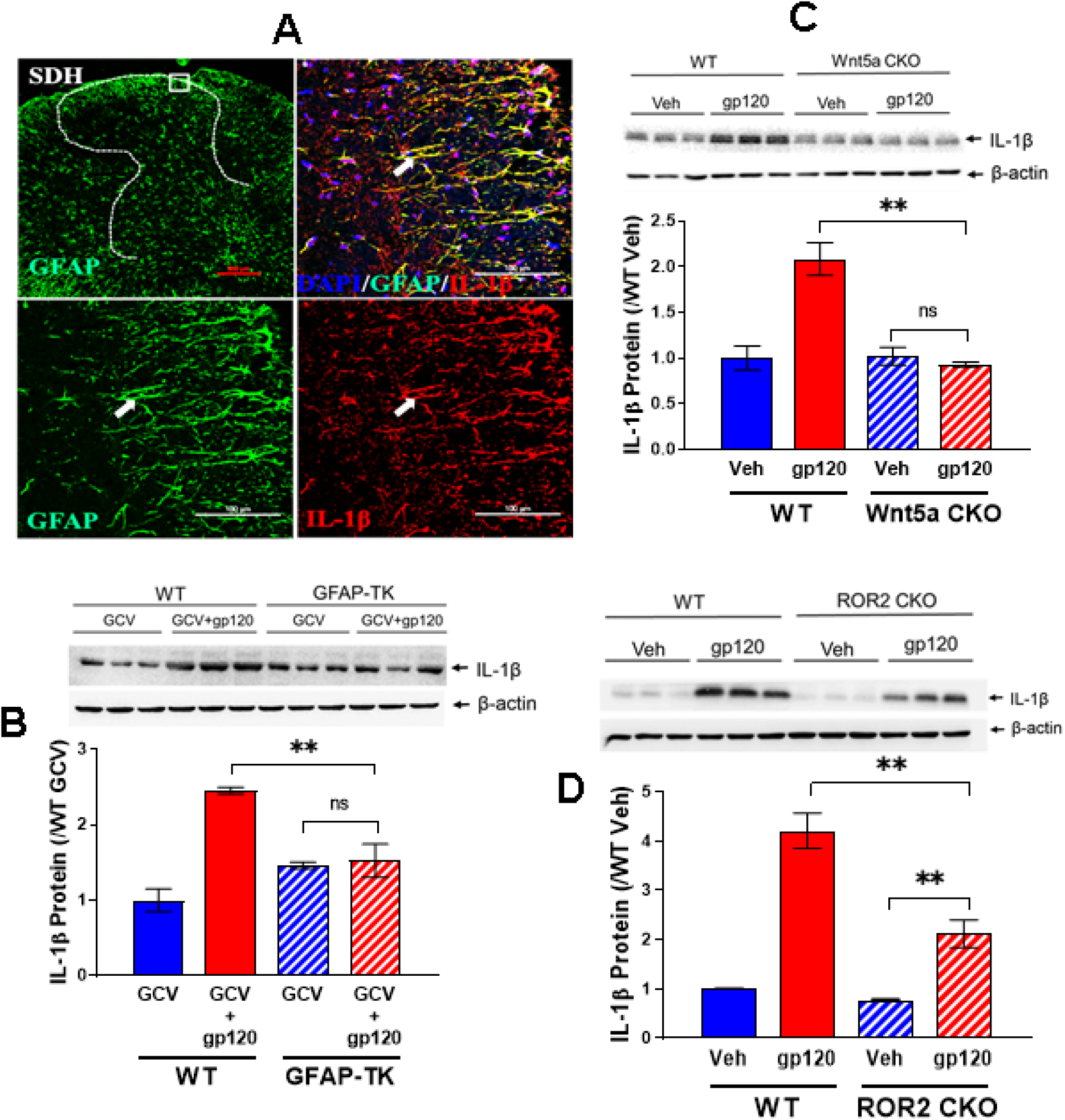
Wnt5a-ROR2 signaling facilitates gp120-induced hyperalgesia and NCP by up-regulating IL-1b in reactive astrocytes. **A**. Immunostaining of GFAP and IL-1β in the SDH from mice treated with gp120 (described in Fig 1A legend). IL-1β signals were mainly observed in astrocytes (arrows). **B**. Immunoblotting analysis showed that ablation of reactive astrocytes in the GFAP-TK mice diminished gp120-induced spinal IL-1β up-regulation. **C**. Neuronal Wnt5a CKO blocked gp120-induced spinal IL-1β up-regulation. Spinal cords were collected for immunoblotting from mice at Day 7 after gp120 administration. **D**. Astrocytic ROR2 CKO partially but significantly inhibited gp120-induced spinal IL-1β up-regulation.

To test the contribution of IL-1β to gp120-induced pain, we used the endogenous IL-1 receptor antagonist IL-1Ra^50^ to block IL-1β signaling. Administration of IL-1Ra (4μg/kg/day in the first 3 days, i.t.) abolished the expression of gp120-induced mechanical hyperalgesia (Fig. 5A), with no detectable effect on baseline mechanical sensitivity (Fig. 5A). These data indicate that IL-1β is crucial for the expression of gp120-induced pain.

**Figure 5.**
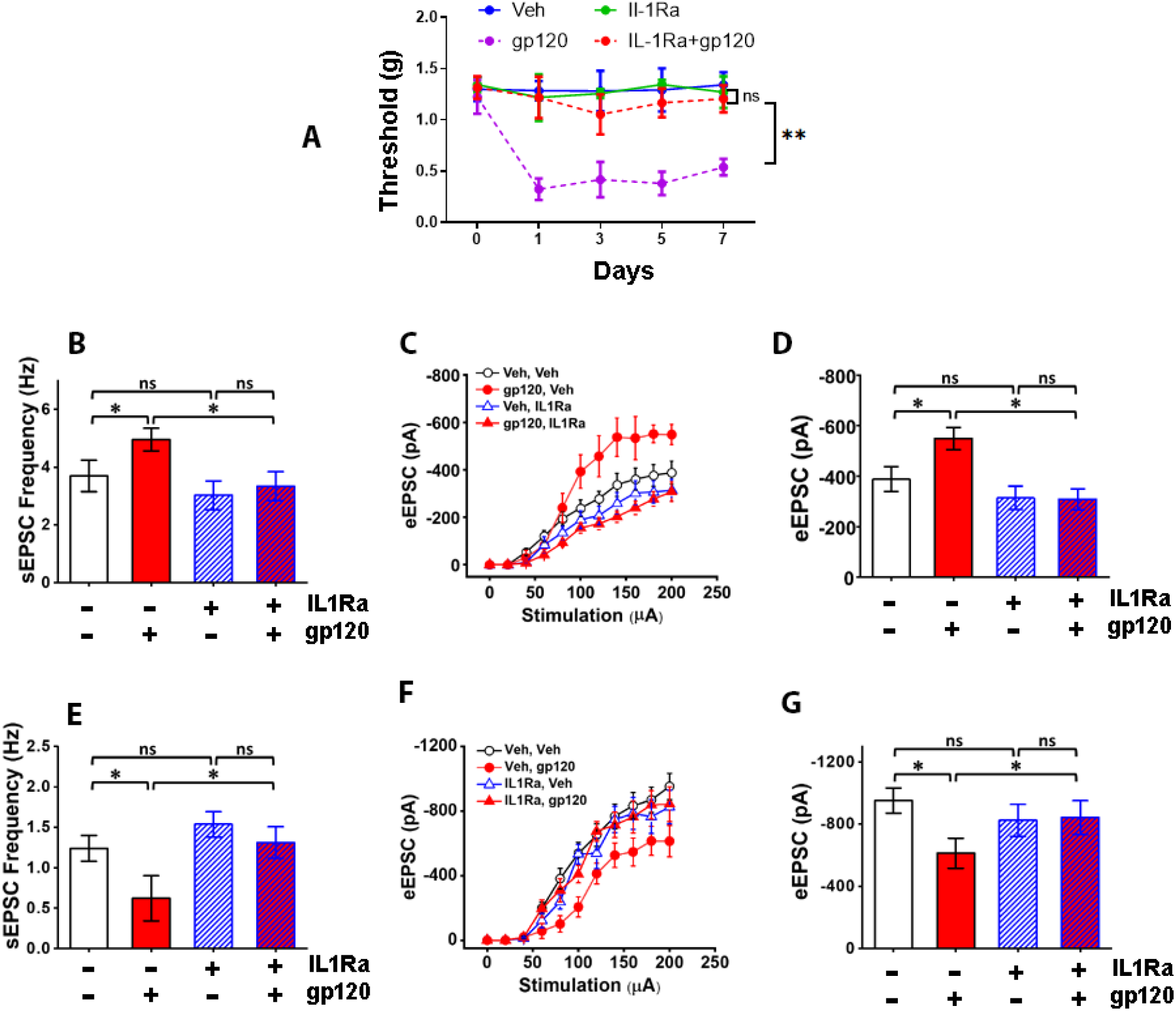
IL-1β is crucial for the expression of gp120-inudced pain. **A**. IL-1β receptor antagonist IL-1Ra (4μg/kg/day, 3 times, i.t.) blocked the expression of mechanical hyperalgesia induced by gp120 (described in Fig 1A legend), measured by von Frey tests (5 mice/group; ns, p>0,05; **, p<0.01). **B.** IL-1Ra impaired gp120-induced increase of sEPSC frequency in non-tonic firing SDH neurons on spinal slices from mice at Day 7 as shown in E. (Veh/Veh: 24/3 (cells/animals); veh/gp120: 50/5; IL-1Ra/Veh: 24/5; IL-1Ra/gp120: 38/3). **C, D**. IL-1Ra diminished gp120-induced increase of eEPSC amplitudes of non-tonic firing neurons in the SDH; H is statistical analysis of eEPSC amplitudes evoked by 200μA stimulation shown in G (Veh/Veh: 24/3 (cells/animals); veh/gp120: 11/3; IL-1Ra/Veh: 19/3; IL-1Ra/gp120: 38/3). **E.** IL-1Ra blocked gp120-induced decrease of sEPSC frequency in SDH inhibitory neurons of GAD67-GFP transgenic mice (Veh/Veh: 30/3 (cells/animals); veh/gp120: 16/3; IL-1Ra/Veh: 35/3; IL-1Ra/gp120: 16/3). **F, G**. IL-1Ra abolished gp120-induced decrease of eEPSC amplitudes of SDH inhibitory neurons; K is statistical analysis of eEPSC amplitudes evoked by 200μA stimulation shown in J (Veh/Veh: 42/4 (cells/animals); veh/gp120: 16/3; IL-1Ra/Veh: 31/3; IL-1Ra/gp120: 16/3). *, p<0.05; ns, p>0,05.

At the neural circuit level, IL-1Ra also impaired the gp120-induced increase in sEPSC frequency (Fig. 5B) and eEPSC amplitude (Fig. 5C, 5D) of non-tonic firing excitatory neurons in the SDH. In addition, using GAD67-GFP transgenic mice^51^, we found that IL-1Ra blocked the gp120-induced decrease in sEPSC frequency (Fig. 5E) and eEPSC amplitude (Fig. 5F, 5G) of GFP-labeled GABAergic inhibitory neurons in the SDH. These results collectively suggest that the expression of NCP induced by gp120 is regulated by IL-1β, which is produced by Wnt5a-ROR2 signaling-activated astrocytes (Fig. 4A-4D).

### Gp120 induces IL-1β via Wnt5a/ROR-regulated MMP2

To understand the mechanism by which gp120 regulates IL-1β, we next tested the role of the inflammasome, a key protein complex controlling IL-1β processing^52^. Surprisingly, the inflammasome inhibitor AC-YVAD-CMK did not affect gp120-induced mechanical hyperalgesia (Sup. Fig. 3A). Furthermore, the inhibitor did not affect gp120-induced increase of active spinal IL-1β neither (Sup. Fig. 3B), although it did block lipopolysaccharides (LPS)-induced IL-1β activation (Sup. Fig. 3C). These data indicate that gp120 might not activate spinal IL-β via inflammasome. Consistent with this notion, we observed that the spinal capase-1 (Cas-1), the inflammasome proteinase that processes pro-IL-1β protein to generate the mature active IL-1β^52^, was not activated by gp120 administration (Sup. Fig. 3D). Prior studies reveal that gp120 is drastically higher in the SDH of pain-positive HIV patients than in the pain-negative HIV patients^2^. However, we did not detect Cas-1 activation in the SDH of pain-positive HIV patients (Sup. Fig. 3E). These findings from both mouse and human patient tissue analyses suggest gp120 does not activate the inflammasome pathway to activate IL-1β in the spinal cord.

Matrix metalloproteinases (MMPs) are implicated in IL-1β processing^53^ and neuropathic pain^54^, and is regulated by gp120^55^. To test the potential roles of MMPs, we used a pan MMP inhibitor GM6001. We observed that GM6001 administration (4μg/kg/day in the first 4 days, i.t.) blocked gp120-induced mechanical hyperalgesia (Fig. 6A), as well as the increase of active IL-1β (Fig. 6B). These data indicated the involvement of MMPs in regulation gp120-induced IL-1β activation. To identify the specific MMP critical for the IL-1β activation induced by gp120, we tested the role of MMP2 and MMP9, both of which have the activity for IL-1β activation^53^. We observed that gp120 up-regulated MMP2 but not MMP9 in the spinal cord (Fig. 6C). We used siRNAs to specifically knock-down MMP2 or MMP9 in the spinal cord (Fig. 6D). We observed that the MMP2 siRNA (3nmol/kg, daily, first 4 days, i.t.) blocked gp120-induced activation of IL-1β activation (Fig. 6E). In contrast, the MMP9 siRNA and scramble siRNA (3nmol/kg, daily, first 4 days, i.t) had no detectable effect (Fig. 6E). Furthermore, we observed that the siRNA targeting MMP2, but not the siRNA targeting MMP9, impaired the maintenance of the gp120-induced mechanical hyperalgesia (Fig. 6F). The early expression phase of the gp120-induced hyperalgesia was not affected by the MMP2 siRNA (Fig. 6F). These data highlight a critical contribution of MMP2 in gp120-induced IL-1β activation during pain pathogenesis.

**Figure 6.**
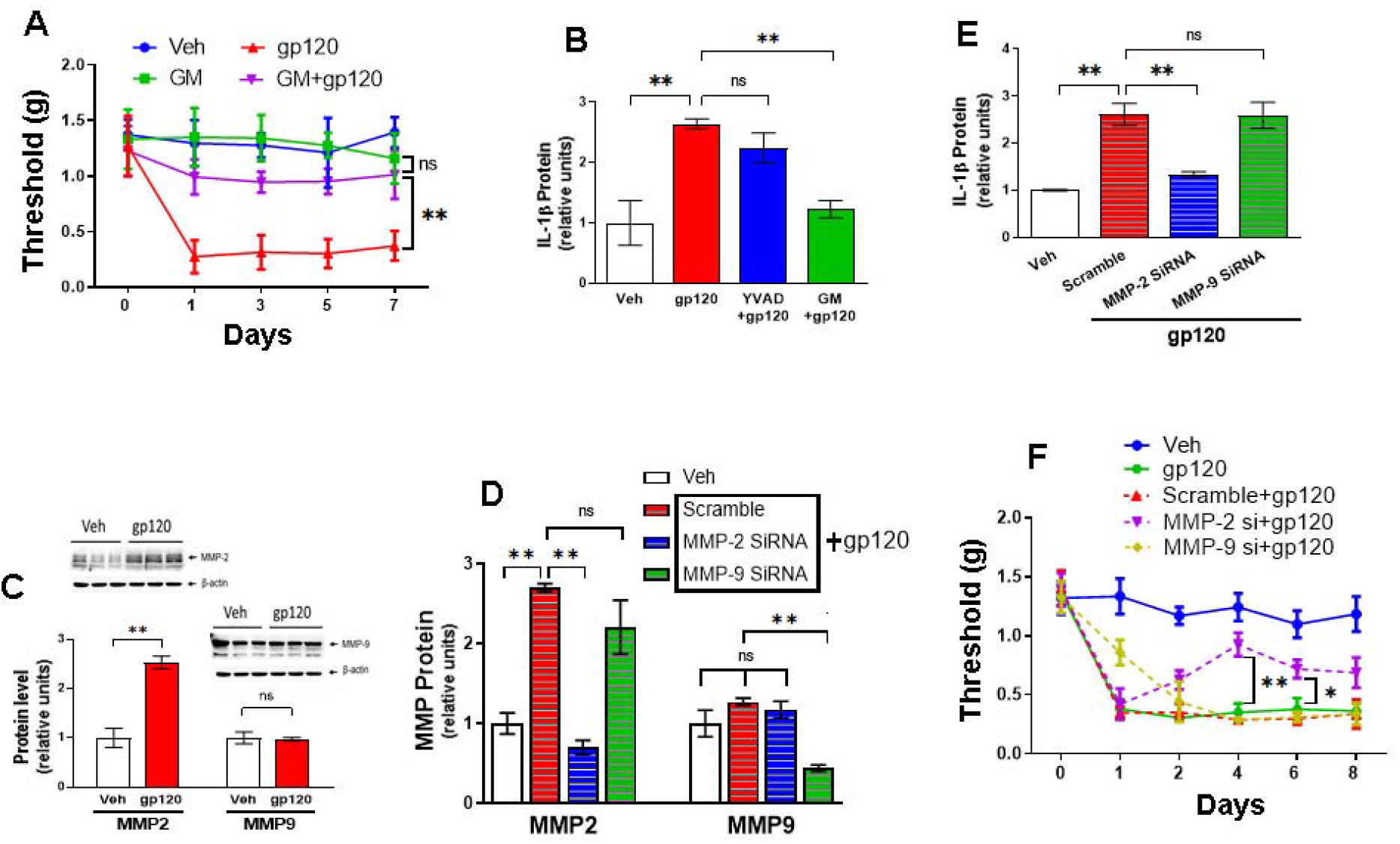
MMP2 is critical for gp120 to induce IL-1β and hyperalgesia. **A**. MMP inhibitor GM6001 (4μg/kg, daily, first 4 days, i.t.) blocked the expression of mechanical hyperalgesia induced by gp120. n=5 mice/group. **, p<0.01: ns, not significant. **B**. Immunoblotting analysis showed that GM6001, but not YVAD, blocked spinal IL-1β increase induced by gp120. Tissues were collected at Day 7. **C**. gp120 induced MMP2 but not MMP9 in the mouse spinal cord (Veh represent control group. n=5 mice/group). **D**. gp120-induced MMP2 protein increase was blocked by MMP2-specific siRNA. Shown are summary data of immunoblotting analysis. n=5 mice/group**. E**. gp120-induced IL-1β was blocked by MMP2-specific siRNA. Shown are summary data of immunoblotting analysis. n=5 mice/group **F**. MMP2 siRNA, but not MMP9 siRNA, impaired the expression of gp120-induced mechanical hyperalgesia (n= 5 mice/group). *, p<0.05; **, p<0.01: ns, not significant.

Finally, we tested the potential role of neuron-to-astrocyte Wnt5a-ROR2 signaling in the regulation of gp120-induced MMP2 upregulation, by determining the effect of neuronal Wnt5a CKO or the astrocytic ROR2 CKO on MMP2 protein levels. We observed that the neuronal Wnt5a CKO abolished the gp120-induced MMP2 increase (Fig. 7A). Again, similar to the effect of the astrogliosis (Fig. 3A) and the IL-1β activation (Fig. 4D), the astrocytic ROR2 CKO not completely but significantly impaired the MMP2 up-regulation induced by gp120 (Fig. 7B). These findings suggest that the neuron-to-astrocyte Wnt5a-ROR2 signaling is crucial for gp120-induced MMP-2 up-regulation.

**Figure 7.**
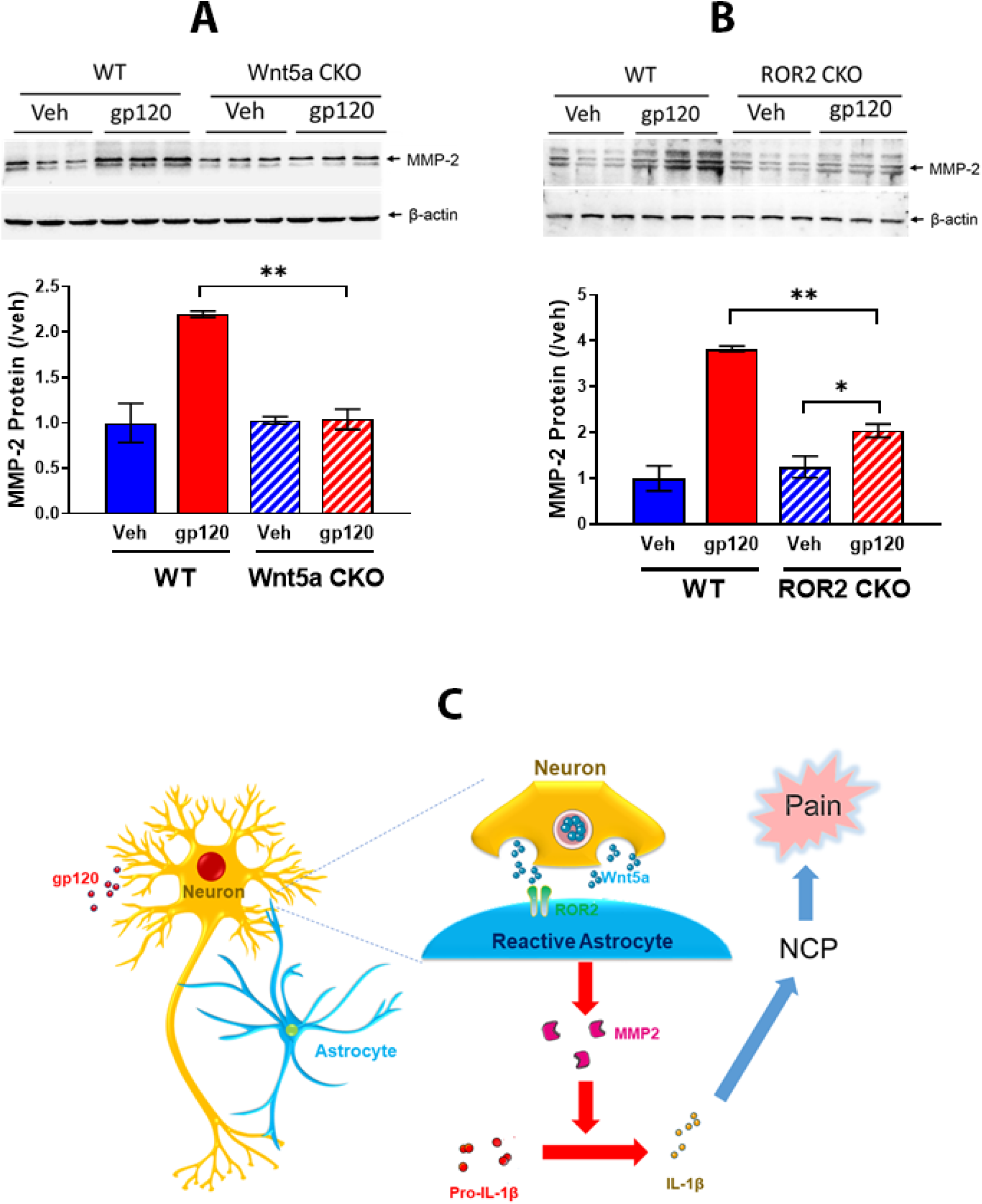
Wnt5a CKO in neurons or ROR2 CKO in astrocytes block gp120-induced MMP2 up-regulation in the SDH. **A**. Neuronal Wnt5a CKO abolished gp120-induced spinal MMP2 up-regulation. n = 5 mice/group. **B**. Astrocytic ROR2 CKO significantly impaired gp120-induced spinal MMP2 protein increase. Tissues were collected at Day 7. n = 5 mice/group; *, p<0.05; **, p<0.01. **C**. A model. The neuron-to-astrocyte Wnt5a-ROR2 signaling pathway regulates the development of gp120-induced pain, via controlling astrogliosis and astrocytic IL-1β activation. IL-1β then induces NCP and pain.

## DISCUSSION

Astrogliosis in the SDH is a neuropathology specifically associated with the manifestation of pain in HIV patients^1^. However, its pathogenic contribution to HIV-associated pain is unclear. The objective of this study is to test the hypothesis that astrogliosis is a critical pathogenic event leading to HIV-associated pain. We provide here complimentary lines of evidence at behavioral, neural circuitry, and molecular levels to validate this hypothesis. We show that HIV-1 gp120 induces mechanical allodynia and pain circuit polarization via astrogliosis. We also elucidate that the astrogliosis is activated by a neuron-to-astrocyte Wnt5a-ROR2 signaling. We further demonstrate that the astrogliosis promotes gp120-induced pain pathogenesis by releasing MMP2-activated IL-1β (Fig. 7C).

### Role of astrogliosis in the development of gp120-induced pain

Astrogliosis is observed in various pain models, but its pathogenic contribution has not been tested conclusively^29^. Prior studies show that astrogliosis is specifically associated with the expression of chronic pain in HIV patients^1^. These findings indicate a critical role of astrogliosis in pain pathogenesis of HIV patients. Results from this study on mouse models of HIV-associated pain provide strong evidence for this hypothesis. We show that ablation of reactive astrocytes in the GFAP-TK mice blocked the expression of mechanical allodynia induced by gp120. In addition, inhibition of astrogliosis by conditional knockout of Wnt5a in neurons or ROR2 in astrocytes also blocks gp120-induced mechanical hyperalgesia. The findings provide complimentary evidence suggesting that astrogliosis plays a crucial role in the pathogenesis of HIV-associated pain. In contrast, microglial ablation only affects partially the early expression of gp120-induced allodynia, indicating a role of microglia during the early stage of the pain pathogenesis. Collectively, these results are consistent with the idea that reactive microglia are mainly involved in the early phase of gp120-induced pain pathogenesis, while reactive astrocytes contribute to both the expression and maintenance of the pain state. Indeed, gp120 appear to induce microglial activation that is more transient than that of astrogliosis^12^. Analyses of postmortem SDH of HIV patients show that the chronic pain state is specific associated with reactive astrocytes but not reactive microglia^1^. The findings of the microglial involvement in the early expression of gp120-induced pain described here indicate that microglia could be transiently activated during the early phase of pain pathogenesis in HIV patients but return to a basal state afterward.

### Neuronal circuitry mechanism of gp120-induced pain

Chronic pain is causally associated with the hyperactivation of pain neuronal circuits in the CNS, commonly known as central sensitization^56^. Pian neural circuits in the SDH of the pain-positive HIV patients and animal model of HIV pain are hyperactivated^1,2^. Diverse molecular and cellular processes have been suggested to contribute to the expression of central sensitization, including dysregulated synapse plasticity and neuroinflammation^56,57^. In this study, we identify a new form of circuitry mechanism potentially underlying the pathogenesis of HIV-associated pain, and we call it neural circuit polarization (NCP). NCP is induced by gp120 in the SDH, and manifested as the increased excitatory synaptic inputs to excitatory neurons and decreased excitatory synaptic inputs to inhibitory neurons. These polarizations are expected to drive hyperactivation of neuronal circuits in the SDH, and thus promote the manifestation of pain. It would be of significance to understand the molecular and cellular basis of NCP in future studies.

### Contribution of astrogliosis to gp120-induced disruption of neural circuit homeostasis

How does astrogliosis promote pathogenesis in HIV-associated pain in neural circuits? Under normal physiological conditions, astrocytes play crucial roles in modulating neuronal activity, by diverse biological processes such as reuptake of extracellular neurotransmitters, release of gliotransmitter and maintenance of cation (e.g., K^+^, Na^+^ and Ca^2+^) and energy metabolism^58,59^. These biological functions could be disturbed when astrocytes are in reactive states, and consequently cause dysregulated neuronal activation in specific circuits. In addition, reactive astrocytes can release inflammatory mediators to stimulate neurons^29^. Astrogliosis and neuronal hyperactivation (e.g. central sensitization) are concurrently observed in the SDH of both pain-positive HIV patients^1^ and the animal models of HIV-associated pain^2^ (and other types of pain such as neuropathic pain). However, a causal relationship between astrogliosis and neuronal hyperactivation has not been established. Our data suggest that astrogliosis is essential for gp120 to induce NCP. With astrocytic ablation, gp120 failed to induce the increase of excitatory inputs on excitatory neurons and the decrease of excitatory inputs on inhibitory neurons. These findings indicate that the astrogliosis mediates the expression of NCP, probably via activating IL-1β (see below). It is of interest that GCV decreases the basal level of excitatory inputs on inhibitory neurons without affecting the excitatory inputs on excitatory neurons. This observation may indicate that astrocytes at the basal state, which could be gently disturbed by GCV, play a critical role in modulating the excitatory inputs to inhibitory neurons.

### Mechanism of gp120-induced astrogliosis

The *in vivo* signals that trigger astrogliosis remain elusive ^29,32^. Multiple signaling pathways have been implicated in astrocyte activation associated with various neurological conditions^60^, including Jak-Stat3 signaling^61^ and the CXCL13-CXCR5 signaling^62^ in pain models. In the current study, we elucidate an intercellular Wnt5a-ROR2 signaling pathway that is crucial for gp120-induced astrogliosis. This finding indicates that gp120 does not activate astrogliosis by directly stimulating astrocytes *in vivo*. Instead, gp120 causes astrogliosis via the Wnt5a-ROR2 intercellular signaling. In support of this notion, gp120 elicits upregulation of spinal Wnt5a and ROR2^41^, which are predominantly expressed in neurons and astrocytes/neurons, respectively^37^. Importantly, Wnt5a is specifically up-regulated in the SDH of pain-positive HIV patients but not of pain-negative patients^38^. We show that, when Wnt5a is specifically deleted in neurons, gp120-induced astrogliosis is blocked, demonstrating that Wnt5a secreted from neurons is essential for the astrogliosis. In addition, when the Wnt5a co-receptor ROR2 is knocked out in astrocytes, the astrogliosis is also blocked. Based on these findings, we propose that gp120 elicits Wnt5a secretion from neurons and the secreted Wnt5a then binds to ROR2 to stimulate astrogliosis (Fig. 7C). This model is consistent with the association of up-regulation of gp120^2^ and Wnt5a^38^ and astrogliosis^1^ in the SDH of the pain-positive HIV patients. Previous studies show that Rho family of GTPase such as RhoA and Rac1 play critical roles in astrocyte activation in various neurological conditions^63,64^. Importantly, Rho GTPases are downstream signaling mediators of the Wnt5a-ROR2 signaling^65^. These findings raise the possibility that Wnt5a-ROR2 signaling is a key regulator of astrogliosis. Astrogliosis occurs in response to various forms of CNS diseases caused by infection, trauma, ischemia, stroke, recurring seizures, chronic pain, or neurodegeneration^30,66^. Gp120 is known to cause hyper-activation of neurons, probably via its co-receptors CXCR4 and/or NMDA receptor^22,67,68^. These diverse insults all cause hyperactivation of neurons in the affected neural circuits. Because Wnt secretion is regulated by neuronal activity^39,40^, we propose that persistent neuronal activation-induced Wnt5a-ROR2 signaling is a common mechanism by which astrogliosis is activated by diverse CNS insults under different neurological conditions, and that this pathway is a potential effective target for developing astrogliosis-targeting therapy for HIV-associated pain and other neurological diseases.

We observe that gp120 requires neuronal Wnt5a to activate astrogliosis. This result indicates that gp120 depends on neurons to activate spinal astrocytes in vivo. gp120 co-receptor CCR5 and CXCR4 were reported to express in astrocytes^69,70^, suggesting stimulation of the co-receptor by gp120 is not sufficient to activate astrogliosis in the spinal cord.

### IL-1β-mediated role of astrogliosis in gp120-induced pain pathogenesis

How does astrogliosis contribute to the development of gp120-induced pain? To answer this important mechanistic question, we tested the hypothesis that astrogliosis promotes gp120-inudced pain pathogenesis via IL-1β. IL-1β is specifically up-regulated in the SDH of pain-positive HIV patients^1^ and induced by gp120 in rodent spinal cords^12,37^, and a key proinflammatory cytokine in pain pathogenesis^71^. We show that spinal administration of IL-1R antagonist blocks gp120-induced mechanical allodynia and NCP. These findings suggest gp120 induces pain and pain neural circuit pathogenesis via IL-1β, and hence indicate the up-regulation of IL-1β in the SDH of pain-positive HIV patients is pathogenically significant^1^.

Both astrogliosis and IL-1β up-regulation are observed in the SDH specifically from HIV patients with chronic pain^1^, raising the possibility that reactive astrocytes produce IL-1β. gp120 induces both spinal astrogliosis and IL-1β up-regulation^2,12^. We observe that IL-1β localization in astrocytes (Fig.4A), like results from studies on other pain models^72,73^. Importantly, ablation of astrogliosis abolished the gp120-induced IL-1β up-regulation (Fig. 4B). These findings provide direct evidence for the idea that gp120-activated astrogliosis produces IL-1β. Further support of this notion comes from our observations that blockage of gp120-induced astrogliosis by neuronal Wnt5a or astroglial ROR2 also impairs gp120-induced IL-1β (Fig.4C, 4D).

We extend our analysis further to elucidate the astrocytic mechanism by which gp120 induces IL-1β. IL-1β is activated from cleavage of pro-IL-1β, commonly thought by inflammasomes^74^, which is also implicated in pain pathogenesis^71^. Surprisingly, we find that complimentary approaches of pharmacological and siRNA inhibition of capase-1 fail to block the IL-1β up-regulation induced by gp120, indicating inflammasomes are dispensable in this process. Consistent with this notion, we find that gp120 administration does not cause inflammasome activation in mouse spinal cords (Sup. Fig. 3D), and that there is no significant activation of inflammations in the SDH from pain-positive HIV patients, where gp120 is drastically higher compared to that from the pain-negative patients^2^. These findings collectively suggest that gp120 does not stimulate inflammasome activation in the spinal cord to activate IL-1β.

We then test the involvement of MMPs, which are implicated in IL-1β activation during the development of neuropathic pain^54^. Our results show that MMP2 but not MMP9 is up-regulated by gp120 (Fig. 6C), and critical for gp120-induced IL-1β activation (Fig. 6E) and mechanical allodynia (Fig. 6F). These data collectively suggest that gp120 induced pain pathogenesis via MMP2-regulated IL-1β.

Finally, we test the role of the Wnt5a-ROR2 signaling in regulation gp120-inudced activation of the MMP2. We find that either the Wnt5a CKO in neurons or the ROR2 CKO in astrocytes blocks gp120-induced increase of the MMP2 (Fig. 7A, 7B). Again, in contrast to the complete abolishing of the MMP2 up-regulation by the Wnt5a CKO, the ROR2 CKO only partially but significantly inhibits the MMP2 up-regulation. This partial inhibitory effect echoes the observations of partial inhibition of astrogliosis and IL-1β up-regulation induced by gp120 (Fig.3A; Fig. 4D). These findings reveal a Wnt5a-ROR2-MMP2-IL-1β axis in gp120-induecd pain pathogenesis.

### Translational potential

Our results reveal that astrogliosis is an important cellular event for the pathogenesis of gp120-induecd pain. This result indicates reactive astrocytes are potential cell target to treat HIV-associated pain. Inhibitory effects of various astroglial inhibitors on non-HIV-associated pain have been reported^75–77^, with the caution of potential off-target effects. It will be of significance to test the selective astroglial inhibitors with HIV pain models.

The Wnt5a-ROR2-MMP2-IL-1β pathway elucidated in this study provides multiple potential molecular targets to for developing drugs to treat HIV-associated pain. In previous studies, we have used a Wnt5a antagonist Box5 on the gp120 pain models, and observed its inhibitory effects on gp120-induced pain^43^. The results from this study provide further support for the translational pursuit on this line of investigation.

Our data suggest that MMP2 is an attractive target, because it is an extracellular proteinase and thus more accessible for drug manipulation. Tremendous efforts have been put to develop selective MMP inhibitors for treatment of cancers and other disease conditions, including pain^78,79^. Testing and improving the specificity of available MMP2 inhibitors could be a productive approach to search for effective medicine for HIV-associated pain.

Our results indicate that IL-1β is another attractive target to treat HIV-associated pain. Pain control by inhibition of IL-1 signaling with IL-1β inhibitors or IL-Ra has been a focal point of drug development of pain medicine for pain associated with various conditions, especially inflammation such as arthritis^80^. Consistent with this theme, we observe that IL-1β is an effector signal from reactive astrocytes that feedbacks to and cause pathogenesis in pain neuronal circuits. Taking the advantage of current available data from preclinical studies and clinical trials, developing IL-1β/IL-1Ra-based drugs could be a productive approach to find an effective medicine for HIV-associated pain.

## Acknowledgement

This work was supported by NIH grants R01NS079166 (SJT), R01DA036165 (SJT), R01NS095747 (SJT) and 1R01DA050530 (SJT, JMC), the Cecil H. and Ida M. Green Distinguished Chair in Neuroscience and Cell Biology (SJT) and U24MH100930 (BBG). All data generated or analyzed during this study are included in this published article and its supplementary information files.

## Author Contributions

Experimental design: SJT; Data collection, analysis and presentation: XL and CB; Experimental Materials: BG; Manuscript preparation: SJT, XL, CB and JMC.

## Materials & Methods

### Animals

Adult male and female C57BL/6 mice (8–10◻weeks), CD11b-DTR (006000; B6. FVB-Tg (ITGAM-DTR/EGFP) 34Lan/J) mice and GFAP-TK (B6.Cg-Tg(Gfap-TK)7.1Mvs/J) mice were obtained from Jackson Laboratory (Bar Harbor, ME, USA). In CD11b-DRT transgenic mice, the expression of diphtheria toxin receptor (DTR) is controlled by the human ITGAM (integrin alpha M) promoter (CD11b). Administration of diphtheria toxin (DT) to the CD11b-DTR transgenic mice induces microglial ablation. In GFAP-TK transgenic mice, herpes simplex virus thymidine kinase (HSV-TK) gene is controlled by the mouse GFAP promoter. Administration of the GFAP-TK mice with ganciclovir (GCV) would specifically ablate proliferating reactive astrocytes cells. Wnt5aflox/flox or Ror2flox/flox mice were generated as previously described (1),(2). Wnt5aflox/flox mice were crossed with B6.Cg-Tg (Syn1-cre) 671Jxm/J mice (Jackson Lab) to produce neuronal Wnt5a conditional knockout mice (CKO; Wnt5aflox/flox/Syn1-cre), and Ror2flox/flox mice were crossed with B6.Cg-Tg(GFAP-cre)77.6Mvs/2J (Jackson Lab) to produce astrocytic Ror2 CKO mice (Ror2flox/flox /GFAP-cre). Animal procedures were performed following protocols approved by the Institutional Animal Care and Use Committee in the University of Texas Medical Branch.

### Materials

HIV-1 gp120Bal recombinant protein (Cat # 4961) were obtained from the NIH AIDS Reagent Program. Ganciclovir sodium (GCV) was purchased from Advanced ChemBlocks (Burlingame, CA, USA). PLX5622-containing rodent diet (each kilogram with 1200mg PLX5622) was purchased from Research Diets (New Brunswick, NJ, USA). Diphtheria toxin (DT), Caspase-1 inhibitor AC-YVAD-CMK (SML0429) and LPS (L3012) were from Sigma-Aldrich (St. Louis, MO, USA). Universal inhibitor for metal matrix proteinases GM6001 (364205) was purchased from Millipore. Flexitube siRNA for MMP2 (GS17390), MMP9 (GS17390) and negative control (s10365635) were purchased from QIAGEN (Germantown, MD, USA). IL-1 receptor antagonist (280-RA) was purchased from R&D Systems Inc. (Minneapolis, MN, USA). Antibodies for immunoblotting: anti-IBa1 (1:1000, Wako: 016-20001), anti-GFAP (1:1000, Millipore: MAB360), anti-β-actin (1:1000, Santa: sc-1616-R), Wnt5a (1:1000, R&D: MAB645-SP), ROR2 (1:1000, LSBio: LS-C334172-20), IL-1β (1:500, Novus: NB600-633), Caspase-1 (1:1000, Adipogen:AG-20B-0042-C100), MMP-2 (1:1000, Santa: sc-10736), MMP-9 (1:1000, PTG:10375-2-AP); Antibodies for immunohistochemistry: anti-IBa1 (1:200, Wako: 016-20001), IL-1β (1:200, Novus: NB600-633), anti-GFAP (1:200, Millipore: MAB360).

### Drug administration

In this study, HIV-1 gp120Bal (4μg/kg) was intrathecally (i.t.) injected to mice daily at the first 2 days and then every other day for total 4 times. DT (0.8μg/kg) was intrathecally injected to gp120-administered CD11b-DRT or wild-type mice daily at the first 2 days. To ablate microglia with PLX5622, C57BL/6 mice were fed with rodent PLX5622-containing diets (1200mg/kg) from 5 days prior to first gp120 injection, and the drug administration continues until the end of experimentation. To ablate reactive astrocytes in the spinal cord, GCV was injected (i.t.; 5mg/kg) to gp120-treated GFAP-TK mice daily at the first 2 days. AC-YVAD-CMK (4nmol/kg), GM6001 (4μg/kg), siRNA MMP-2 (3nmol/kg) and siRNA MMP-9 (3nmol/kg) were intrathecally injected to gp120-administered wild-type C57BL/6 mice daily for first 4 consecutive days. IL-1 receptor antagonist (IL-Ra; 4μg/kg) was intrathecally injected to gp120-treated wild-type C57BL/6 mice daily for first 3 consecutive days. In the LPS experiment, LPS (40μg/kg) and AC-YVAD-CMK (4nmol/kg) were intrathecally injected to wild-type C57BL/6 mice sequentially, with a 30 seconds interval. Mice treated with drug vehicle were used as controls.

### Measurement of mechanical nociception by Von Frey test

Mechanical nociceptive sensitivity was measured by Von Frey test as described (3). Before testing session, mice were habituated by being placed on a wire mesh platform (1mm diameter wire with a 4×4 mm square grid pattern) for 40 minutes daily for 3 days. Six calibrated Von Frey filaments ranging from 0.07 to 2 g (Stoelting, Wood Dale, IL) were used. When testing session began, the 0.4 gram filament was applied from below the wire mesh to the central area of plantar surface of a mouse left hind paw when it was resting and standing on four paws. A complete hind paw lift from the platform was recorded as a withdrawal response (marked as “X”) and then a less size filament was applied after at least 1 minute and when the mouse was stationary. If the specific filament application did not cause a withdrawal response (marked as “O”), a next larger size filament would apply to repeat the process until got enough “O” and “X” records required by Dixon “up and down paradigm” method. After testing the threshold of the left hind paw, the repeated procedure was carried out on the right hind paw 5 minutes later and the average of withdrawal threshold of the left and right hind paw will be considered as the final value of withdrawal threshold. The baseline withdrawal thresholds of each of the hind paws were determined at least 2 hour prior to the first time of drug administration (Day 0). All tests were blindly conducted by an experimenter who did not know details of the mouse strain and drug administration.

### Western blotting analysis

Mice were anesthetized with 3% isoflurane and sacrificed. L4-L6 lumbar spinal cord segments were collected. Spinal tissues were homogenized in RIPA lysis buffer (1% Nonidet P-40, 50◻mM Tris–HCl, pH 7.4, 150◻mM NaCl, 0.5% sodium deoxycholate, , 1◻mM EDTA, pH 8.0) with a protease inhibitor cocktail (Sigma). Protein concentration of the lysates was determined by BCA Protein Assay Kit (Thermo). Equal amounts of protein (2μg) were separated on 12% SDS-PAGE and transferred to polyvinylidene fluoride membrane. The membrane was blocked with 5% nonfat milk in TBST buffer (Tris-buffered saline with 0.1% Tween 20) for 1 hour at room temperature, and then incubated with primary antibodies in TBST buffer overnight at 4℃. After brief washes (10 Min × 3 times) with TBST buffer in room temperature (25℃), membranes were incubated with HRP-conjugated secondary antibody. Enhanced Chemiluminescence kit (Pierce) were used to visualize protein bands, and the intensity of protein bands was quantified with NIH Imagej software. β-actin was used as a loading control

### Immunofluorescence staining

Mice were anesthetized with 3% isoflurane and transcardially perfused first with 20◻ml of 0.01M PBS (0.14 M NaCl, 0.0027 M KCl, 0.010 M PO43), followed by 30◻ml of 4% paraformaldehyde (PFA) in 0.01◻M phosphate buffer in ice. The L4 and L5 lumbar spinal cord segments were dissected and fixed in 4% PFA solution for 12◻hours at 4°C. The fixed tissues were dehydrated with 30% sucrose solution in PBS for 24 hours at 4°C, and then embedded and frozen in optimal cutting temperature medium (Tissue-Tek). Tissues were sectioned (15 μm) on a cryostat (Leica CM 1900), and sections were mounted onto Superfrost Plus microscope slides (Fisher Scientific, Waltham, MA, USA). For fluorescence staining, sections were first incubated in blocking solution (5% BSA, 0.3% Triton X-100 in 0.01◻M PBS) for 1◻h at room temperature and then with primary antibodies diluted in blocking solution overnight. Sections were washed with 0.01M PBS (10 Min × 3 times) and incubated in FITC or texas red-conjugated secondary antibody (1:200, Jackson ImmunoResearch Laboratories) in dilution buffer (1% BSA and 0.3% Triton X-100 in 0.01M PBS). After washes in 0.01M PBS (10 Min × 3 times), sections were mounted in mounting medium with DAPI (Sigma). IgG from the same animal sources as the primary antibodies was used as negative controls. Images were collected with a laser scanning confocal microscope (model A1, Nikon S).

### Patch-clamp recording of dorsal horn neurons in ex vivo spinal cord slices

Spinal cord slices were prepared as previously described (4). Briefly, the spinal cord was sliced transversely at a thickness of 350 μm using a vibratome (Leica VT1200S, Buffalo Grove, IL) in cold (~4oC) NMDG (N-methyl-D-glucamine) solution (in mM: 93 NMDG, 2.5 KCl, 1.2 NaH2PO4, 30 NaHCO3, 20 HEPES, 25 glucose, 5 sodium ascorbate, 2 thiourea, 3 sodium pyruvate, 10 MgSO4 and 0.5 CaCl2, pH 7.4), saturated with 95% O2 and 5% CO2. Whole-cell recordings were made on random neurons in lamina II in artificial cerebrospinal fluid (ACSF in mM: 124 NaCl, 2.5 KCl, 1.2 NaH2PO4, 24 NaHCO3, 5 HEPES, 12.5 glucose, 2 MgSO4, and 2 CaCl2, pH 7.4) using Multiclamp 700B amplifier, DigiDATA, and pClamp software (version 10.6. Molecular Device, Sunnyvale, CA) at a 10 kHz sampling rate and a 2 kHz filtering rate. The patch-pipettes (4–8 MΩ) were filled with internal solution (in mM: 120 K-gluconate, 10 KCl, 2Mg-ATP, 0.5 Na-GTP, 0.5 EGTA, 20 HEPES, and 10 phosphocreatine, pH 7.3). After making whole-cell recording configuration, the spontaneous excitatory postsynaptic currents (sEPSC) were recorded for 60 second at −65 mV in ACSF. EPSCs were evoked by focal electrical stimulation in the vicinity of recorded neurons with a metal bipolar electrode (MicroProbes, Gaithersburg, MD). Test pulses were given for 0.5 ms at 5 second intervals and stimulation intensities ranging from 20 μA to 200 μA (20 μA step). Recording was made only when the evoked EPSCs (eEPSCs) were monosynaptic, based on eEPSC waveforms with a short latency, single peak, and stable responses to repeated stimuli. All recordings showing polysynaptic response were disregarded. The neurons were characterized by their action potential (AP)-firing pattern upon depolarizing current injections in the current clamp mode (5).

### Statistical analysis

Statistical analysis was conducted with Prism 5 (GraphPad) software. Data were represented as mean◻±◻SEM from three independent experiments. One-way ANOVA was used for immunoblotting data and two-way ANOVA with a Bonferroni post hoc test was performed for analyzing animal pain behavior. Electrophysiological data are expressed as mean◻±◻standard error of the mean (SEM) with n, the number of cells and N, the number of animals. The sEPSC frequency and eEPSC amplitude were analyzed off-line using Clampfit software (version 11, Molecular Devices, CA). sEPSC events were detected using the template event detection method. These lectrophysiological data were analyzed using two-way analysis of variance followed by Holm–Sidak multiple comparison tests. In all tests, p◻<◻0.05 was considered significant. The Origin (version PRO 2020, OriginLab, Northampton, MA) was used to analyze the data.

## Supplementary Figures

**Supplemental Figure 1.**
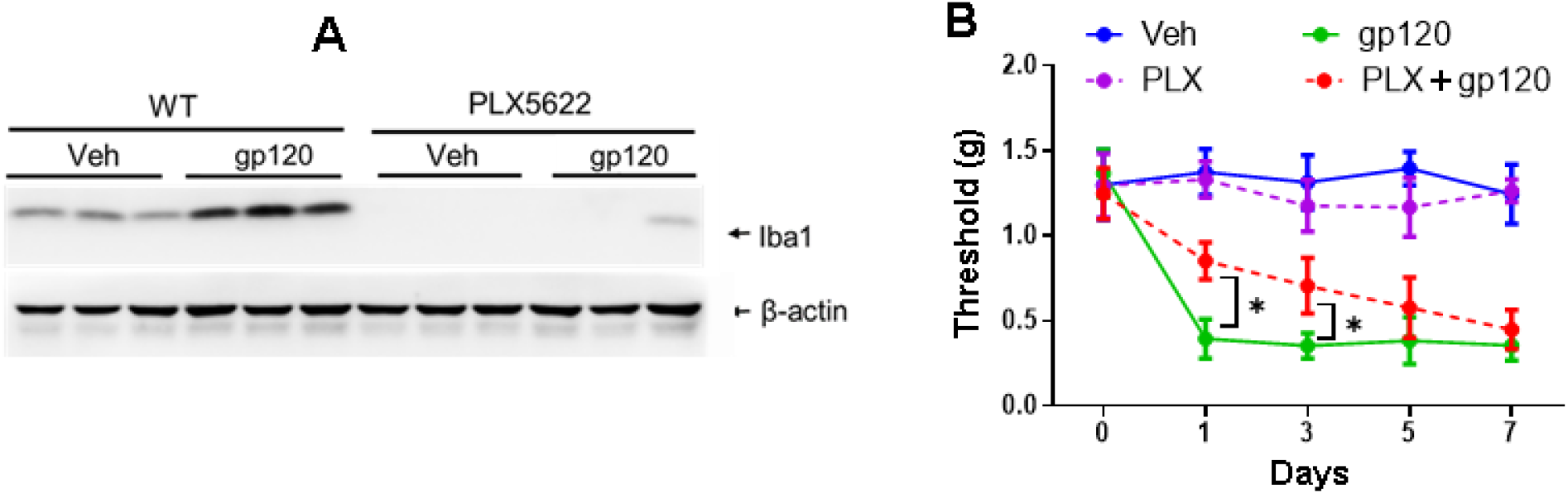
Microglial ablation only affects the early phase of gp120-inducd hyperalgesia. **A**. Immunoblotting analysis of spinal microglia showed the activation of microglia by gp120 (described in Fig 1A legend) and the depletion of microglia by PLX5622 (1200mg/kg, from 5 days prior to first administration of gp120 until the end of experimentation). Tissues were collected at Day 7; **B**. Effect of microglial depletion on the expression of gp120-induced mechanical hyperalgesia measured by von Frey tests (Administration of gp120 and PLX5622 were described in Sup. Fig 1A). 5mice/group, *, p<0.05

**Supplemental Figure 2.**
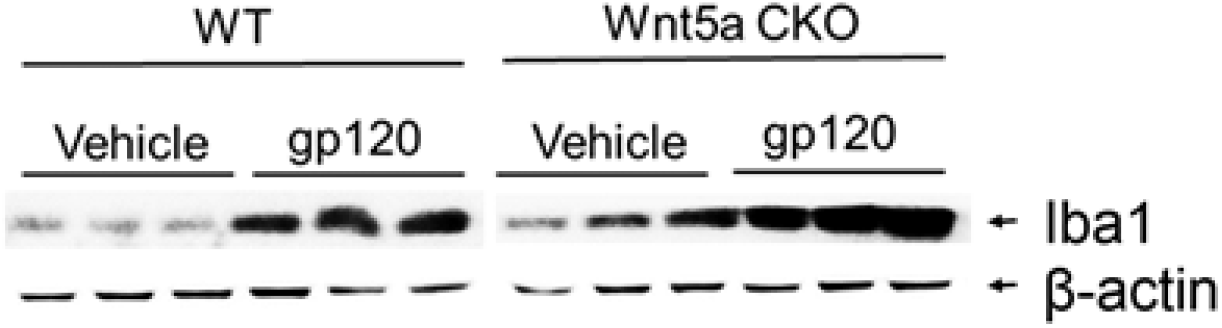
Wnt5a CKO in neurons does not affect gp120-induced microglial activation. Immunoblotting analysis of spinal Iba1 from mice at Day 7 after gp120 administration (described in Fig 1A legend).

**Supplemental Figure 3.**
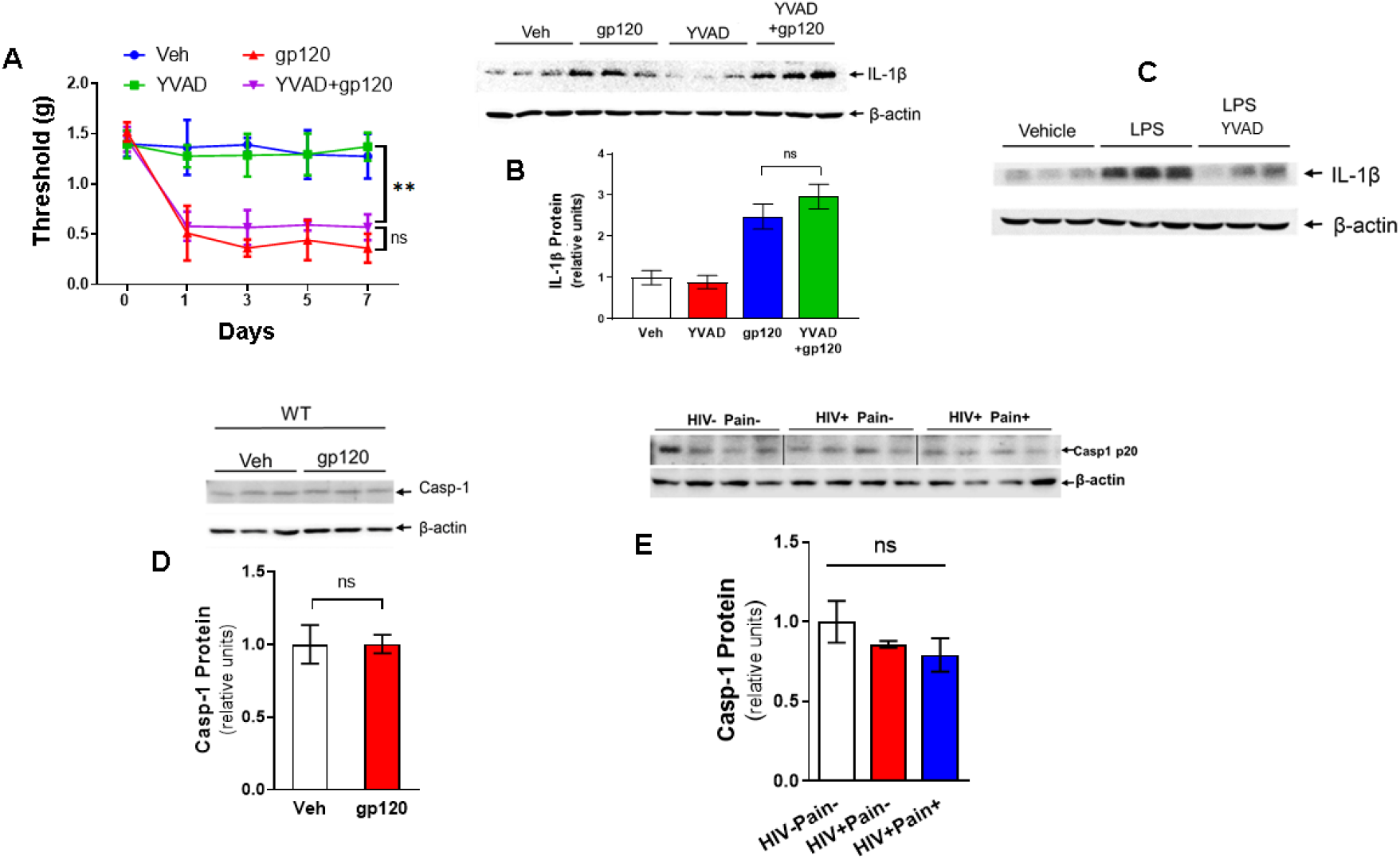
Gp120 induces IL-1β via an inflammasome-independent mechanism. **A**. Casp-1 inhibitor YVAD had no detectable effect on gp120-induced mechanical hyperalgesia measured by von Frey tests. n=5 mice/group. **, p<0.01: ns, not significant **B**. YVAD (4nmol/kg, daily, first 4 days, i.t.) had not apparent effect on the spinal IL-1β up-regulation induced by gp120 (described in Fig 1A legend). **C**. YVAD blocked spinal IL-1β up-regulation induced by LPS (40μg/kg, one time, 30 second prior to administration of YVAD, i.t.). Spinal cords were collected at 24 hours after treatment. **D**. gp120 (described in Fig 1A legend) did not induce casp-1 activation in the mouse spinal cord. Tissues were collected at day 7. **E**. Immunoblotting analysis of caspase-1 (casp-1) in the postmortem spinal tissues indicated that inflammasome was similarly activated in the cohorts of HIV patients (with or without chronic pain) and non-HIV patients.

